# Effect of choline-phosphate cytidylyltransferase gene (*pcyt-1*) expression on departure of pine wood nematode, *Bursaphelenchus xylophilus* (Nematoda: Aphelenchoididae), from *Monochamus alternatus* (Coleoptera:Cerambycidae)

**DOI:** 10.1101/2020.02.22.943852

**Authors:** Yang Wang, Fengmao Chen

## Abstract

In order to study the causes of pine wood nematodes (PWN), *Bursaphelenchus xylophilus* departure from its vector beetle, *Monochamus alternatus*, we collected PWNs which were extracted from the newly emerged *M. alternatus* and the 7d after emergence beetles. The total RNAs of the two groups of PWNs were extracted and transcriptomes sequencing were performed, and the genes expression differences between the two groups of PWN were analyzed. It was found that expression of choline-phosphate cytidylyltransferase gene (*pcyt-1*) markedly up-regulated. After inhibition of *pcyt-1* expression by RNA interference, the rate of lipid degradation in PWN decreased significantly, and the motility of PWN also decreased significantly. Analysis identified that phosphatidylcholine can promote the emulsification and degradation of lipid droplets in PWN, which provide energy for PWN departure from *M. alternatus*. The up-regulation of gene *pcyt-1* is an important internal trigger for PWN departure from the beetles.

## 1 Introduction

Pine wilt disease (PWD) constitutes one of the most serious conifer diseases worldwide, affecting *Pinus* spp. from the Far East forestlands (Japan, China, Korea) [1, 2], and from Europe (Portugal and Spain) [3-6]. PWD is caused by the pine wood nematode (PWN), *Bursaphelenchus xylophilus* (Steiner & Buhrer) Nickle, with PWN transmission being dependent on a vector insect, such as *Monochamus alternatus* Hope, the main vector in East Asia [7, 8]. The feeding period of *M. alternatus* is an important stage in the life cycle of beetles, as well as a key step in PWN transmission [9, 10]. After emergence, *M. alternatus* feed on healthy pine trees, a process accompanied by the spread of PWN.

PWN has two developmental forms in its life cycle, namely the propagative and dispersive forms. Under favorable conditions, PWN molt into their propagative form, and then reproduce rapidly [11, 12]. However, under unfavorable conditions, e.g. high temperature, starvation, or high population density, PWN (as propagative second-stage juveniles) will molt to produce the dispersive third-stage juveniles, which will aggregate around the pupal chamber of the beetles [13, 14]. The dispersal third-stage juveniles molt to produce fourth-stage juveniles, which enter the tracheal system of the vector as the beetles emerge [13-16], the fourth-stage juveniles to be transmitted to the healthy pine trees through wounds caused by the vector [17-20].

There is no unified view on the mechanism of PWN departure from vector. However, it has been found that some volatile chemicals play an important role in this process of PWN exit from the vector. It was found that the monoterpenes released from healthy pine trees, such as *β* - myrcene and α-pinene, had the strongest attraction to PWN, these chemicals playing an important role in the process of PWN departure from vector and the invasion of healthy host trees, as well as in the movement of PWN within pine trees [21, 22]. Ishkawa et al. found that *β* - myrcene on the agar plate held obvious attraction for dispersive third-stage juveniles [23]. Enda et al. dried *Pinus* twigs at 70°C to remove volatile substances, then treated twigs with *β*-myrcene and fed them to the beetles. It was found that these twigs greatly promoted the departure of PWN from *M. alternatus*[24]. Stamps et al. showed that various volatile pine chemicals did not significantly affect PWN departure from the vector and the departure of PWN is a spontaneous act [25]. Aikawai et al. fed beetles were fed with fresh pine twigs and with twigs treated with high temperature (121°C, 40min, to remove volatiles from the pine twigs), respectively, and found that the pine twigs without volatile chemicals were more likely to cause PWN departure from vector, while twigs with volatiles had inhibitory effects [26].

The departure of PWN may depend on the nematode’s own factors, or it may be environmental factors, or a combination of the two. Stamps et al. suggested that the level of lipids in PWN may be the switch that determines whether they depart from the vector [20]. When PWN had low neutral storage lipid content, departure is induced by the pine volatile *β*-myrcene, whereas, when PWN had a high content of neutral lipids, they continued to be retained within the beetle [20]. Previous studies shown that intrinsic factors play an important role in regulating the PWN departure, whereas *β*-myrcene is a directional signal that attracts PWN escape from the vector insects [25]. Yang et al. there was no difference in the time of PWN departure from *M. alternatus* which between fed after starvation and fed directly, and the motility of PWN carried by the beetles one week after emergence being greater than the PWN carried by newly emerged (almost resting) beetles [27]. It was speculated that PWN departure was related to the motility, and the increased motility required energy which might come from the degradation of neutral lipids [20, 27].

The emulsification of lipid droplets can increase the contact area between lipases and lipids, which greatly promote lipids degradation[28]. In the present study, transcriptome sequencing was performed on the PWN which carried by new emerged *M. alternatus* and beetles one week after their emergence. Inhibition of expression of the *pcyt-1* gene, encoding choline-phosphate cytidylyltransferase (EC 2.7.7.15), was carried out by RNA interference (RNAi), and the effect on lipid degradation and motility of PWNs was determined.

## 2 Materials and methods

### 2.1 PWN collection and RNA extraction

In April 2019, dead specimens *Pinus massoniana* trees, infested by *M. alternatus* larvae and PWN, were collected at Bocun Forest Farm, Huangshan City, Anhui Province in China. The trees were cut into logs and maintained in outdoor insect cages (**Fig. 1A**). *M. alternatus* were collected daily (every 8 h) during the period of adult beetles emergence. The collected *M. alternatus* were divided into two groups, and one of which PWN was extracted immediately with Baermann funnel method[29], whereas another was fed (with fresh pine twigs) for seven days (**Fig. 1B**), before the PWN was extracted. Extraction of total RNAs from the two groups (each group was represented by three independent biological replicates) of PWN by Trizol method [30].

**Fig. 1.**
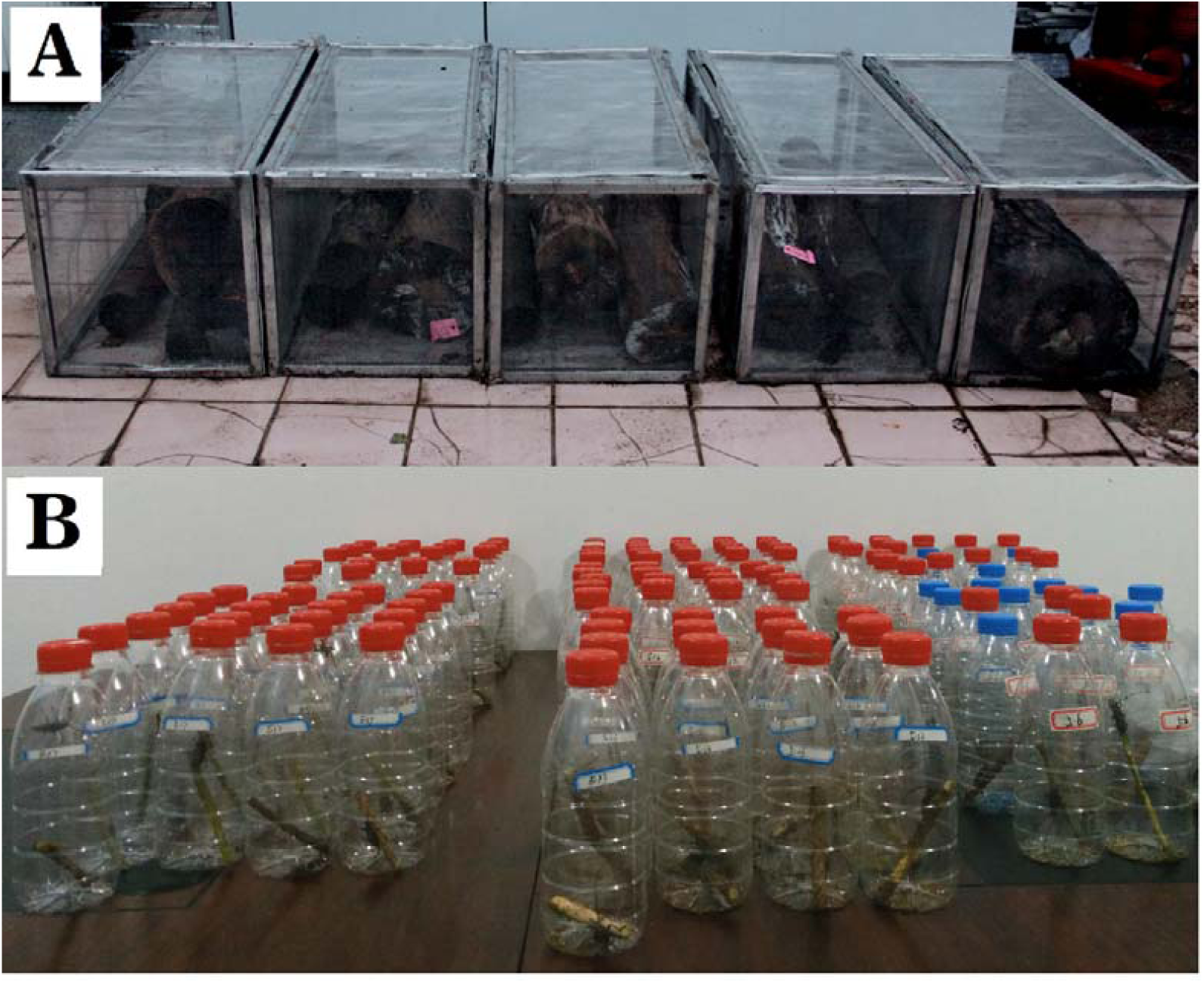
Collection and feeding of *Monochamus alternatus*. A: Pine trees were cut into logs and maintained in outdoor insect cages, *Monochamus alternatus* were collected daily (every 8 h) during the period of adult beetle emergence. B: The collected *Monochamus alternatus* were put into small bottles and fed with fresh twigs of pine.

### 2.2 Transcriptome sequencing and analysis (Data provided by Biomarker Technologies, Beijing, China)

#### RNA quantification and qualification

RNA degradation and contamination was monitored on 1% agarose gels. RNA purity was checked using the NanoPhotometer® spectrophotometer (IMPLEN, CA, USA). RNA concentration was measured using Qubit® RNA Assay Kit in Qubit®2.0 Flurometer (Life Technologies, CA, USA). RNA integrity was assessed using the RNA Nano 6000 Assay Kit of the Agilent Bioanalyzer 2100 system (Agilent Technologies, CA, USA).

#### Library preparation for Transcriptome sequencing

A total amount of 1 μg RNA per sample was used as input material for the RNA sample preparations. Sequencing libraries were generated using NEBNext®Ultra™ RNA Library Prep Kit for Illumina® (NEB, USA) following manufacturer’s recommendations and index codes were added to attribute sequences to each sample. Briefly, mRNA was purified from total RNA using poly-T oligo-attached magnetic beads. Fragmentation was carried out using divalent cations under elevated temperature in NEBNext First Strand Synthesis Reaction Buffer (5X). First strand cDNA was synthesized using random hexamer primer and M-MuLV Reverse Transcriptase (RNase H-). Second strand cDNA synthesis was subsequently performed using DNA Polymerase I and RNase H. Remaining overhangs were converted into blunt ends via exonuclease/polymerase activities. After adenylation of 3’ ends of DNA fragments, NEBNext Adaptor with hairpin loop structure were ligated to prepare for hybridization. In order to select cDNA fragments of preferentially 200-250 bp in length, the library fragments were purified with AMPure XP system (Beckman Coulter, Beverly, USA). Then 3 μl USER Enzyme (NEB, USA) was used with size-selected, adaptor-ligated cDNA at 37°C for 15 min followed by 5 min at 95°C before PCR. Then PCR was performed with Phusion High-Fidelity DNA polymerase, Universal PCR primers and Index (X) Primer. At last, PCR products were purified (AMPure XP system) and library quality was assessed on the Agilent Bioanalyzer 2100 system.

#### Clustering and sequencing

The clustering of the index-coded samples was performed on a cBot Cluster Generation System using TruSeq PE Cluster Kit v4-cBot-HS (Illumia) according to the manufacturer’s instructions. After cluster generation, the library preparations were sequenced on an Illumina Hiseq x-ten platform at Biomarker Technologies (Beijing, China) and paired-end reads were generated.

#### Quality control

Raw data (raw reads) of fastq format were firstly processed through in-house perl scripts. In this step, clean data(clean reads) were obtained by removing reads containing adapter, reads containing ploy-N and low quality reads from raw data. At the same time, Q20, Q30, GC-content and sequence duplication level of the clean data were calculated. All the downstream analyses were based on clean data with high quality.

#### Comparative analysis, and Gene functional annotation

The adaptor sequences and low-quality sequence reads were removed from the data sets. Raw sequences were transformed into clean reads after data processing. These clean reads were then mapped to the reference genome sequence. Only reads with a perfect match or one mismatch were further analyzed and annotated based on the reference genome.

Gene function was annotated based on the following databases: Nr (NCBI non-redundant protein sequences); Nt (NCBI non-redundant nucleotide sequences); Pfam (Protein family); KOG/COG (Clusters of Orthologous Groups of proteins); Swiss-Prot (A manually annotated and reviewed protein sequence database); KO (KEGG Ortholog database); GO (Gene Ontology).

#### Differential expression analysis

Differential expression analysis of two conditions/groups was performed using the DESeq R package (1.10.1). DESeq provide statistical routines for determining differential expression in digital gene expression data using a model based on the negative binomial distribution. The resulting P values were adjusted using the Benjamini and Hochberg’s approach for controlling the false discovery rate. Genes with an adjusted P-value <0.05 found by DESeq were assigned as differentially expressed.

#### KEGG pathway enrichment analysis

KEGG is a database resource for understanding high-level functions and utilities of the biological system, such as the cell, the organism and the ecosystem, from molecular-level information, especially large-scale molecular datasets generated by genome sequencing and other high-throughput experimental technologies (http://www.genome.jp/kegg/). We used KOBAS software to test the statistical enrichment of differential expression genes in KEGG pathways.

### 2.3 RNA interference (RNAi) and qPCR (quantitative real-time PCR) of PWN (pine wood nematodes) after pcyt-1 interference

To interfere with the choline-phosphate cytidylyltransferase gene (***pcyt-1***) (DNA sequence: **Supplemental 1**), the interfering gene fragment was 5’-GGTGGACTTTATCGCTCAT-3’, and negative control sequence is 5’-GGTGG**G**CTTTA**A**CGCTCAT-3’, and designed the antisense strand [31]. To prepare the double-stranded RNA (dsRNA), which was digested by ribozyme to obtain the small interfering RNA (siRNA) (dissolved in 20 μl of RNAase-free water), the *in vitro* Transcription T7 Kit (for siRNA Synthesis) (Takara Bio Inc., Shiga, Japan) was used.

The PWN was cultured using the *Botrytis cinerea* medium [32], as shown in **Fig. 2.** After 48 h of culture, PWN was extracted (for 2 h) by the Baermann funnel method [29]. Extracted PWN was soaked in 0.05% (w/v) streptomycin sulfate for 10 min and washed three times with diethyl pyrocarbonate (DEPC)-treated water. PWNs (10000 individuals) were soaked in the siRNA solution for 48 h (**Fig. 2E**), and then washed three times with RNAase-free water. The total RNAs (one preparation per replicate per group) were extracted by the Trizol method [30]. Synthesis of cDNA was carried out using HiScript^®^ II Q RT SuperMIX for Qpcr (+gDNA wiper) (Vazyme, Nanjing, China). qPCR was carried out with SYBR^®^ Green I chimeric fluorescence, using ChamQ™ SYBR^®^ qPCR Master Mix (Low ROX Primixed) (Vazyme, Nanjing, China). Then q-PCR validation was performed using the 7500 Real-Time PCR System (SeqGen, Inc. Torrance, CA, USA) and analyzed differences in gene expression. The reference gene primer was F: 5’-GCAACACGGAGTTCGTTGTA-3’, R: 5’-GTATCGTCACCAACTGGGAT-3’. Target primer: F: 5’-AGGCCTACAACAACTCAGCC-3’, R: 5’-TCAACTGATTCGCGTGTCCA-3’.

**Fig. 2.**
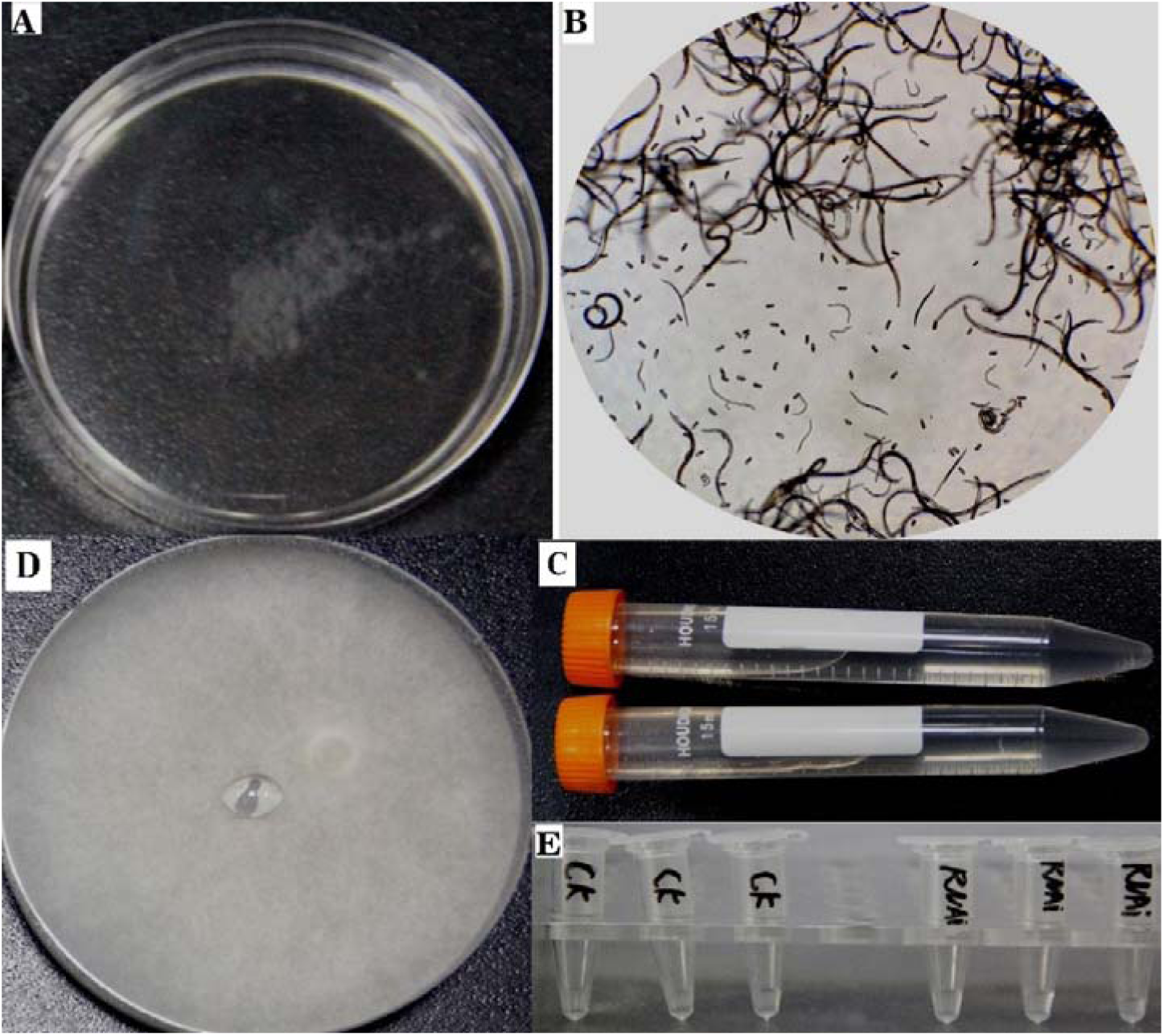
PWN culture and RNAi. A: PWN suspension was placed in a dish, and the eggs adhering to the bottom of the dish were collected every 6 h. B: Eggs adhering to the bottom of the dish. C: The collected PWN eggs were placed in a Eppendorf tube and incubated at 25 ° C for 36 h. All the eggs hatched into second-stage PWN. D: The second-stage PWN juvenile suspension was inoculated into the freshly grown *Botrytis cinerea* medium and cultured for 48 h (when the majority of nematodes were 4th stage juvenile). **Notes**: 1. Freshly cultured *Botrytis cinerea* medium was used. Aging medium is not conducive to PWN growth. 2. Experiment was performed using PWN of the same stage. Mixed ages can cause large errors. 3. The time taken to extract PWN should not be too long, with 2-4 h being suitable.

### 2.4 Comparison of the difference in motility and lipids content of PWN after *pcyt-1* RNAi

After 24 h of RNAi (at which time, there was a great difference in motility between the two PWN groups), the differences of PWN motility were observed, with videos being taken under a microscope (Zeiss Axio lmager.M2, Carl Zeiss, Gottingen, Germany). After 72 h of RNAi (at which time, the lipids contents of the two PWN groups were markedly different.), the lipids of PWN were stained with oil red O [33]. PWN was washed into a 1.5ml eppendorf tube with M9 buffer (standing for 10min at 4°C, discarding the supernatant and repeating it once). The lipids of PWN were fixed with 4% paraformaldehyde at 4°C for 30 minutes, then frozen in a - 80°C refrigerator for 15 minutes, then thawed rapidly in 43°C water bath. The sample was centrifuged for 1 min with 2000 × g, and the supernatant was discarded. Wash three times with 1 × phosphate-buffered saline (PBS) (pH 7.2-7.4), after which the PBS was discarded, and soaked in 1% Triton X-100 and isopropanol oil red O saturated solution (ratio 2:3) for 30min. Wash three times with PBS (1x), useing 60% isopropanol to differentiate clearly under the microscope, and photos were taken (Zeiss Axio lmager. M2, Carl Zeiss, Gottingen, Germany).

## 3 Results

### 3.1 Comparison of transcriptome results

The expression level of choline-phosphate cytidylyltransferase gene (***pcyt-1***) of PWN carried by *M. alternatus*, 7 d after of emergence, was significantly up-regulated (log2 FC=9.5) relative to PWN from newly emerged beetles, as shown in supplementary material 2 (Ko00564, EC:2.7.7.15).

### 3.2 q-PCR gene experession studies

After RNAi to *pcyt-1* was carried out in PWN, q-PCR was used for verification of transcript abundance and the results are shown in Fig. 3, indicating that, after RNAi, the expression level of ***pcyt-1*** was significantly down-regulated.

**Fig. 3.**
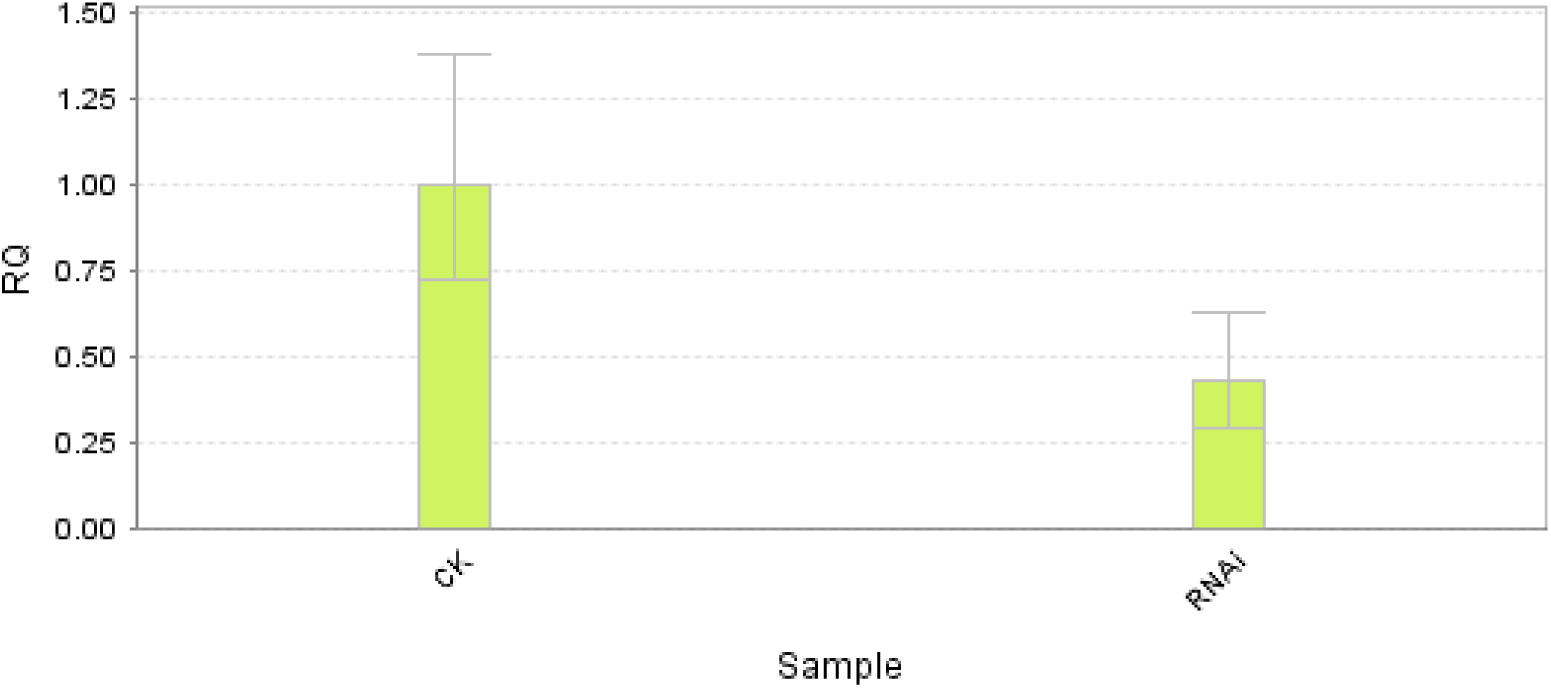
Relative expression of *pcyt-1* in control pine wood nematodes (PWN) (CK) and PWN exposed to RNA interference (RNAi).

### 3.3 Observation on the motility and lipids content of PWN

After RNAi, the motility of PWN decreased significantly, as shown in supplement material 3 (videos 1 and 2). After RNAi, the lipids content of PWN was significantly higher than that in the control nematodes (Fig. 2).

## 4 Discussion

PWN did not departure immediately after the emergence of *M. alternatus*, but started to departure after a period of time. There have been reports that PWN start to departure from *M. alternatus* at different times, such as 7–12 d after the emergence of beetles [34], at 3–5 d after adult beetle emergence [35], at 7 d after emergence [36], at 5 d after emergence [27, 37], at 10 d after emergence [38]. During the period between the emergence of *M. alternatus* and the start of the departure of PWN, the physiological characteristics of PWN may have changed, or it may be continuously stimulated by certain signaling hemicals. Different studies have different conclusions on the causes of PWN departure from *M. alternatus*.

Ishkawa et al. examined the attractant effect of volatile chemicals from *Pinus densiflora* towards the dispersive PWN that leave *M. alternatus*, and showed that β-myrcene played an important role in the transmigration of the PWN from the vector to the pine tree and for the movement of the PWN inside the pine wood [23]. To clarify the effect of the volatiles of *Pinus densiflora* on the departure of PWN from *M. alternatus*, Aikawa et al. fed *M. alternatus* with normal twigs and with twigs without volatiles after heating, and consequently suggested that PWN had a trait of spontaneous departure from *M. alternatus* [26]. Stamps et al. found that lipid content of PWN gradually decreased with the prolongation after the emergence of *M. alternatus*, and suggested that the concentration of neutral storage lipids was correlated with the date at which PWN started to leave their vector [18, 20].

In order to study the effect of feeding behavior of *M. alternatus* on PWN departure. Wang et al. found that there was no significant difference in the start time of PWN departure from *M. alternatus* direct feeding and feeding after starvation after emergence[27]. However, it was found that PWN carried by the newly emerged beetles was less motility than the PWN carried by the beetles at 7 d after emergence, and believed that the increase of motility is an important cause of PWN departure[27]. The increased motility requires a lot of energy (in the form of ATP). The degradation of neutral storage (NS) lipid in PWN can provide energy[20]. Therefore, it is concluded that the degradation of lipid may be an important endogenous cause of PWN departure. In the present study, we also found that the lipid content of PWN after RNAi of *pcyt-1* was markedly higher than that of the control, whereas the motility of PWN in the RNAi-treated group was lower than the control. This indicates that lipid degradation is a necessary pre-condition for PWN departure from *M. alternatus*.

Not all PWN that carried by *M. alternatus* can leave the beetles. About 70% of PWN failing to leave the vector [34]. Aikawa et al. showed that the percentage of PWN departure form vector was higher in thick tracheae than in thin tracheae [39]. And considered that PWN in the thin trachea cannot get enough oxygen, so they can’t produce enough energy (ATP) to power their exit, and eventually cause they staying in the trachea for a long time [39]. After the emergence of *M. alternatus*, the energy resource (represented by the limited lipid store) of PWN carried by beetles is gradually exhausted [20]; at this time, even if getting sufficient oxygen, it would not generate enough energy to drive PWN to move. Wang et al. showed that greater motility was necessary for PWN to departure from the vector [27]. It is vital for PWN departure to get sufficient oxygen to degrade lipids and produce enough energy (ATP) over a limited period, enabling the PWN to achieve high motility.

Lipid droplets (triacylglycerol) are the main energy store in PWN, with lipid being insoluble in water whereas the enzymes that digest the lipid are water-soluble, with lipid digestion occurring only at the lipid–water interface [28]. The emulsification of lipid can increase the lipid-water interface, which can greatly accelerate its digestion. Phosphatidylcholine (PC) has hydrophilic and hydrophobic groups, making it a powerful “detergent” conducive to emulsifying lipid and promoting lipid catabolism [28, 40-42]. Choline-phosphate cytidylyltransferase (CCT) has high specificity and is the rate-limiting enzyme of phosphatidylcholine synthesis, controlling the synthesis of phosphatidylcholine [43-45]. In the present study, the results of transcriptome sequencing revealed that gene ***pcyt-1*** expression level was significantly up-regulated in PWN in beetles 7 d after emergence, relative to those extracted from beetles immediately after emergence ((supplementary 2: ko00564, EC: 2.7.7.15). Following RNAi with the gene ***pcyt-1***, the level of lipid catabolism decreased significantly (**Fig. 4**) and the motility of PWN decreased significantly (supplementary 3). The results showed that PC emulsification of lipid in PWN is an important route of lipid emulsification, with the level of ***pcyt-1*** expression having a significant effect on nematode departure.

**Figure 4.**
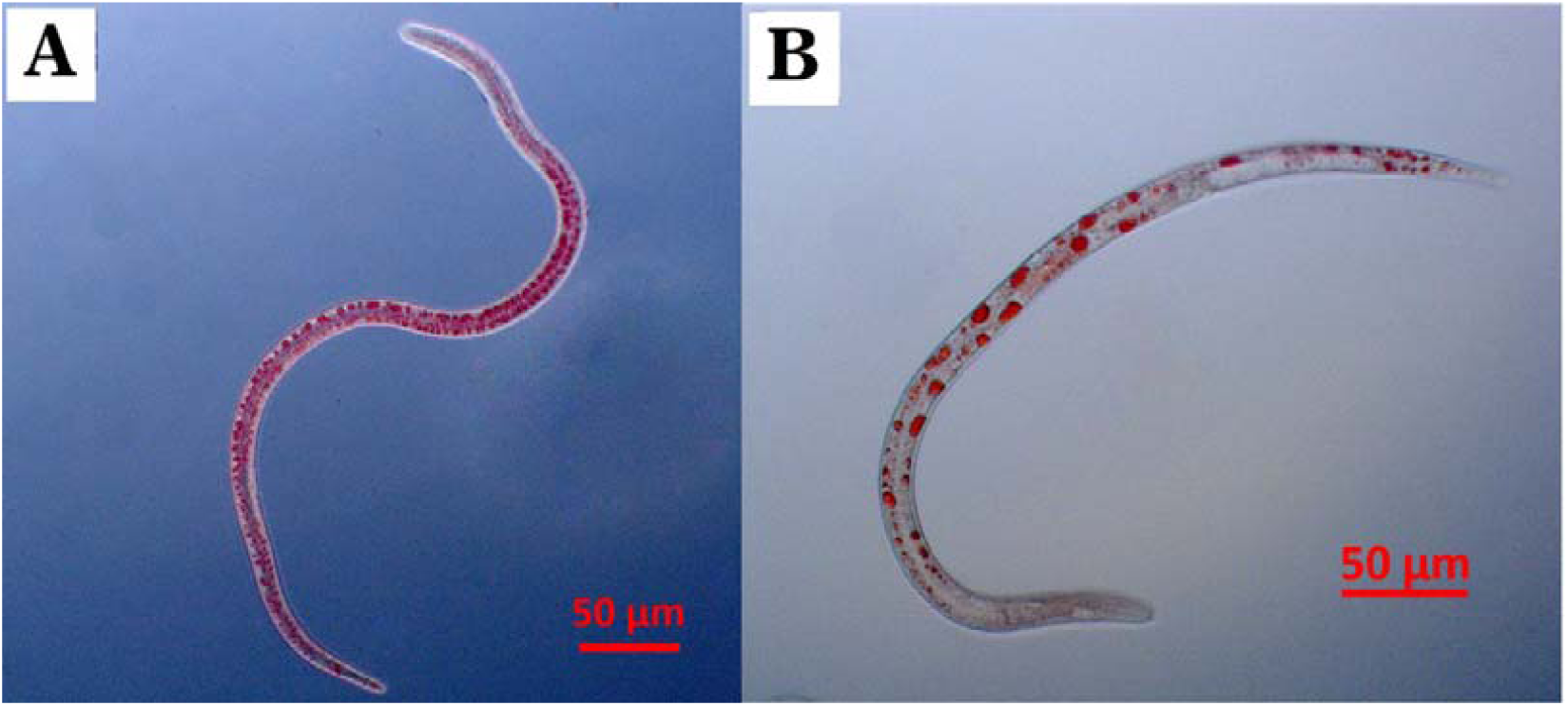
Pine wood nematode lipid content as shown by staining with Oil Red O in the presence and absence of RNA interference (RNAi) for *pcyt-1*. Red spots are lipid drops. A: Nematodes after RNAi for 72 h. B: The control nematodes after culture 72 h. There were still many lipid droplets in the nematodes of the experimental group, while the number of lipid droplets in the control group was markedly less.

In the present study, the results showed that no eggs were laid in the experimental group (after RNAi), unlike the situation in the control group (supplementary 3). This indicates that the expression level of gene ***pcyt-1*** affects the development of PWN, suggesting that PWN in the trachea of *M. alternatus* may have some development, which may promote the departure of PWN from the vector. However, this development process also needs amount of energy. Therefore, the degradation of Lipids is still an important factor for PWN departure from *M. alternauts*.

## Supporting information

Supplementary gene sequence

Supplementary kegg pathway

