## Supplementary figures and images for "Effect of choline-phosphate cytidylyltransferase gene (*pcyt-1*) expression on departure of pine wood nematode, *Bursaphelenchus xylophilus* (Nematoda: Aphelenchoididae), from *Monochamus alternatus* (Coleoptera:Cerambycidae)"

### ko00564.png

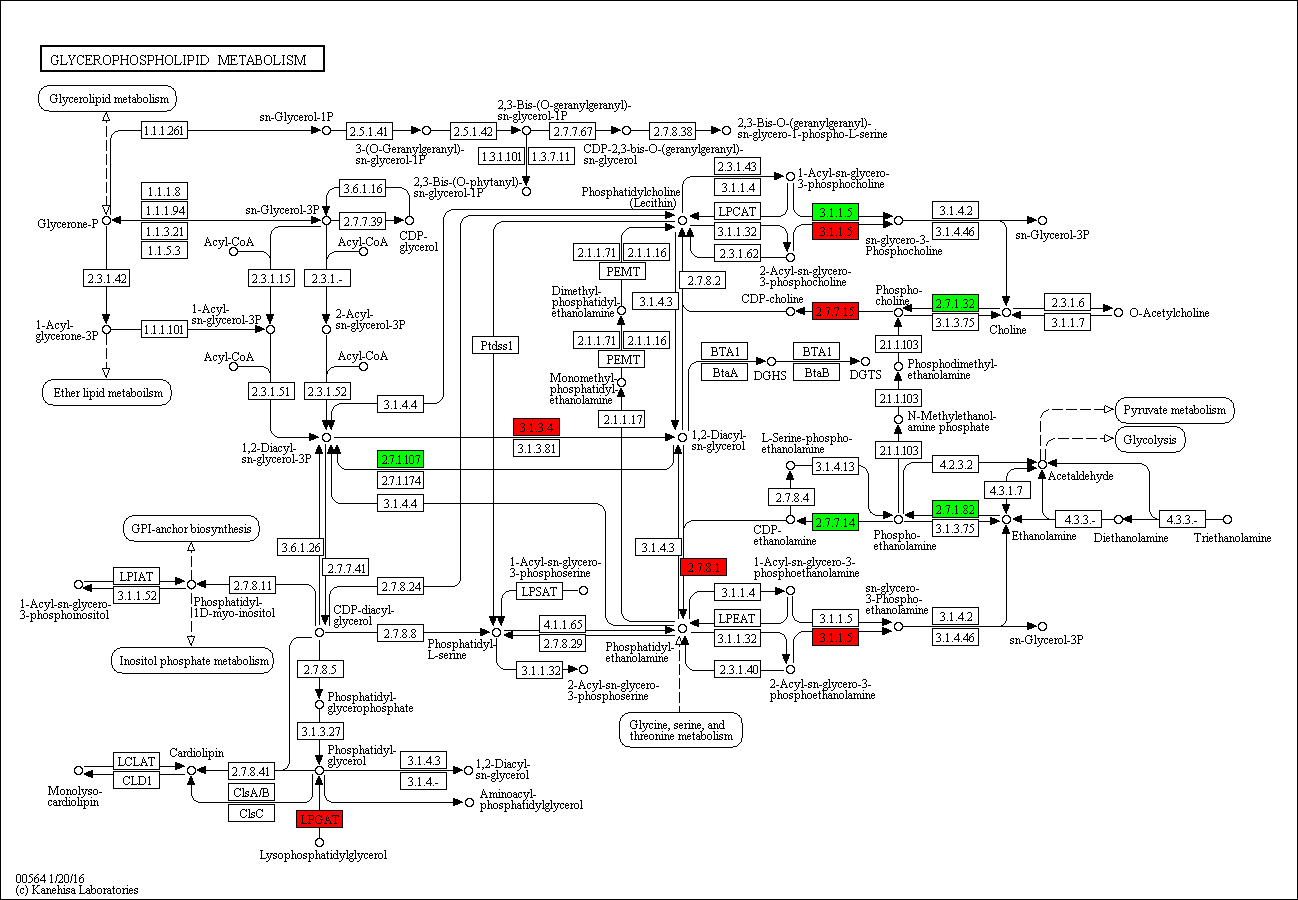
